# The impact of spatial and temporal dimensions of disturbances on ecosystem stability

**DOI:** 10.1101/429100

**Authors:** Yuval R. Zelnik, Jean-François Arnoldi, Michel Loreau

## Abstract

Ecosystems constantly face disturbances which vary in their spatial and temporal features, yet little is known on how these features affect ecosystem recovery and persistence, i.e. ecosystem stability. We address this issue by considering three ecosystem models with different local dynamics, and ask how their stability properties depend on the spatial and temporal properties of disturbances. We measure the spatial dimension of disturbances by their spatial extent while controlling for their overall strength, and their temporal dimension by the average frequency of random disturbance events. Our models show that the return to equilibrium following a disturbance depends strongly on the disturbance’s extent, due to rescue effects mediated by dispersal. We then reveal a direct relation between the temporal variability caused by repeated disturbances and the recovery from an isolated disturbance event. Although this could suggest a trivial dependency of ecosystem response on disturbance frequency, we find that this is true only up to a frequency threshold, which depends on both the disturbance spatial features and the ecosystem dynamics. Beyond this threshold the response changes qualitatively, displaying spatial clusters of disturbed regions, causing an increase in variability, and even the loss of persistence for ecosystems with alternative stable states. Thus, spanning the spatial dimension of disturbances can allow probing of the underlying dynamics of the ecosystem. Furthermore, considering spatial and temporal dimensions of disturbances in conjunction is necessary to predict the type of ecosystem responses that can have dramatic consequences, such as regime shifts.

## I. Introduction

Understanding the stability of ecosystems, i.e. their ability to recover and persist in the face of natural and anthropogenic disturbances, is of fundamental importance to ecology and conservation [15, 17, 22]. Ecosystems are spatially extended, comprised of multiple interacting communities in different locations, and therefore an important factor in understanding their stability is their spatial structure [14, 25, 33]. However, while the influence of space on properties such as biodiversity and food web structure has been intensely investigated [5, 16, 18, 21], basic questions regarding spatial stability remain open. In particular, despite the fact that most disturbances (e.g. fires, pest outbreak, pollution runoff) are strongly heterogeneous in space, the impact of their spatial structure on stability is largely unknown. The natural counterpart of this spatial dimension of disturbances is their temporal dimension, e.g. their timespan or the frequency of their occurrence. Taken together, these dimensions span a vast space of possible disturbances that ecosystems can face. This, in part, explains why reaching a clear understanding of ecosystem stability has proven to be an extremely challenging endeavor.

Research on ecosystem stability has a long history in ecology, and numerous studies have investigated how various properties of disturbances affect ecosystem responses. The importance of spatial properties of disturbances, in particular, has been assessed by a few studies of regeneration dynamics under recurrent, spatially structured disturbances [10, 20, 32]. These studies introduced the concept of landscape equilibrium and demonstrated how the spatial and temporal scales of disturbances can generate different stability patterns. A point not explicitly addressed in these studies, however, is the importance of rescue dynamics occurring at a regional scale when local recovery processes are too slow or fail altogether. This can occur in sufficiently connected ecosystems, following high-intensity disturbances [9, 10]. In fact, recovery from a disturbance is a consequence of both local and regional processes. Local processes lead to recovery due to dynamics that are internal to local communities, while regional processes lead to recovery by bringing in individuals from neighboring communities via dispersal [13, 30]. These two processes mediate the large-scale system response to a disturbance, and their respective parts in this response is bound to strongly depend on the spatial connectivity of the system and, importantly, on the spatial structure of disturbances.

Recent work has made this relationship more explicit, by defining three distinct regimes of a system’s recovery from a single spatially heterogenous disturbance: Isolated, Rescue and Mixing [36]. If a system is highly connected due to strong dispersal of organisms, then it is in the Mixing Regime, and the system’s behavior at large scales is essentially an extended version of a local system [7]. At the other extreme, if dispersal is low and hence each site acts separately with its own local dynamics, then the system is in the Isolated Regime, and its large-scale behavior is an aggregation of many independent small systems [28, 35]. In between these two extremes is the Rescue Regime, where systems with intermediate connectivity show large-scale rescue dynamics due to the interaction between limited dispersal and the system’s behavior at the local scale [6, 24, 34]. While spatial structure of both system and disturbance plays no role in the Mixing regime, for weaker dispersal it does: in both the Isolated Regime and the Rescue Regime the spatial structure of the disturbance has significant effects as it can initiate qualitatively different responses that involve both local and regional processes [36]. We will therefore consider systems with intermediate dispersal, and focus on the effect of the spatial structure of disturbances as well as their temporal properties.

Quantifying the impact of disturbances amounts to defining relevant stability measures. If the disturbance is an isolated event, a natural measure to consider is the return time to the unperturbed state [17, 22]. On the other hand, in a regime of repeated disturbances (e.g. climatic events), measures of temporal variability are commonly used [29]. In the presence of alternative stable states, those repeated disturbances can cause a regime shift from one state to another. Here the stability measure of interest is typically persistence, i.e. the probability that a system will remain in a desired state [11, 26]. Importantly, these stability measures reflect not only the spatial and temporal properties of the disturbance, but also the dynamical features of the perturbed ecosystem. Exploring this interplay is the focus of our study, which we will address by considering three spatial ecosystem models with increasing nonlinear local dynamics, ranging from logistic growth to bistability. Under various perturbation scenarios we will measure their stability using return time, variability and persistence.

We begin by looking at the ecosystem’s recovery following a single disturbance, and show that changing the spatial structure of the disturbance reveals two basic recovery trajectories: isolated and rescue. Isolated recovery trajectories reflect the local resilience of the system, while rescue trajectories involve spatial processes, and their dominance signals the failure of local processes. We thus argue that the relationship between spatial structure and recovery contains substantial information about the local dynamics of the system, both close and far from equilibrium. We continue by exploring the temporal axis of disturbances, and demonstrate a direct link between return time (following an isolated disturbance event) and temporal variability (under a regime of repeated disturbances). We find that for low disturbance frequency patterns of variability do not contain additional information in comparison to the patterns of return time. However, past a frequency threshold (which depends on the system’s internal dynamics) the variability patterns change. As we will argue, this signals the onset of a new dynamical regime driven by disturbances, which can lead to a regime shift, and hence a loss of persistence.

Our work demonstrates that the spatial dimension of disturbances can be used to reveal information on the ecosystem’s internal behavior. Furthermore, the conjunction of the spatial and temporal properties of disturbances may lead to unforeseen dynamical responses, with drastic ecological consequences.

## II. Methods

### A. Models

We assume for simplicity that the local community dynamics can be described by a single state variable *N* that describes the ecosystem’s local biomass density. We study the dynamics in multiple locations in space using partial differential equations. We define three different models that differ in their local dynamics but that describe identical dispersal across space with linear diffusion. In all models the local biomass may reach a carrying capacity *K*, so that *N* = *K* (the populated state) is a stable steady state in of all three models. An additional solution exists for *N* = 0 (the bare state), with its stability properties differing among models.

The first and simplest model (LG) describes local logistic growth and dispersal: 
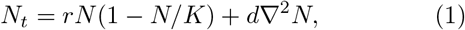
where *N_t_* is the change in time of the local biomass and ∇^2^*N* is the second derivative in space of *N* (a diffusion term). Here *r* is the intrinsic per capita growth rate, while the rate of spread by dispersal is governed by *d*. In this model the bare state *N* = 0 is an unstable solution.

The second model (AE) describes species dynamics with an Allee effect, so that low biomass densities are not viable: 
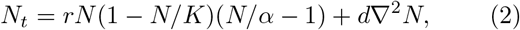
where *α* is the viability threshold, i.e. the minimal amount of biomass *N* that is necessary to maintain a viable density. This model has two alternative stable steady states (*N* = 0, *N* = *K*) and a single unstable steady state (*N* =α), and we assume that 0 *< α < K*.

Finally, the third model (SR) shows an intermediate nonlinearity between the first and second models. This model is useful in clarifying the distinction between strong nonlinearity and bistability, which will be discussed in the Results section. Its main feature is that while there is only one stable equilibrium at *N* = *K*, far from this equilibrium the return rate is very slow compared with the return rate close to equilibrium. This could model succession dynamics, for which the recovery following strong disturbances (e.g. clearcutting) is very slow, as it involves the successive colonization by different species, and not simply the regrowth of the disturbed species. The model is: 
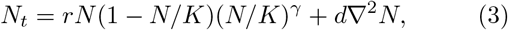
where *γ* controls the nonlinearity of the dynamics, such that at high values of *γ* local recovery is very slow following high-intensity disturbances. The other parameters are the same as in the previous models.

By rescaling time, space and biomass, we can effectively reduce the parameter space of the models, and set *r* = 1, *d* = 1 and *K* = 1. Our results thus hold for any values of these three parameters. We set *α* = 0.4 to make sure that the AE model recovers from a single disturbance (see next subsection), and *γ* = 4 to make sure the return time far from equilibrium of the SR model is sufficiently slow. We focus on one-dimensional systems as they are simpler to analyze, but the qualitative results hold for other types of spatial structure such as two-dimensional systems (see appendix D). We use a system size of *L* = 500, which is large enough to allow for the spatial dynamics to manifest itself (so that the system is not in the Mixing Regime [36]), with periodic boundary conditions. For a clearer illustration, in Fig. 3 and Fig. A2 we show snapshots of a two-dimensional system of size 200 × 200.

### B. The spatial dimension of disturbances

We define a disturbance as a change in the state variable that is forced on the ecosystem. We consider a pulse disturbance occurring at a given time, with its full effect being applied at that time. This assumption is appropriate for the many types of disturbances that are faster than the dynamics of the ecosystem, and it allows a simpler analysis. We choose a disturbance that removes biomass (reduces *N*), so that a disturbance of strength *s* will reduce the overall biomass of the ecosystem by *sK*(but any negative values of *N* will be set to 0 for consistency). Once a disturbance takes place, the ecosystem may recover to its original state, or a regime shift can occur if the ecosystem is bistable. We are interested here in stability and recovery dynamics, and therefore focus on parameter values for which a single disturbance cannot lead to a regime shift.

Since a disturbance need not occur uniformly across space, we vary the spatial extent of the disturbance σ while keeping its overall strength *s* constant. A disturbance is performed by choosing its locus, and removing some biomass in a domain of size σ centered around the locus. We can vary the spatial extent from σ = 1 for a uniform disturbance across space, to σ = *s* for a localized disturbance.

To measure recovery we use the return time *T* defined as the time needed for the ecosystem to recover 90% of the biomass lost to the disturbance. While the choice of a threshold is arbitrary, its specific value has no significant effect on the results as long as it is not too close to either 0% or 100% (which roughly correspond to reactivity and asymptotic resilience, respectively [2]). By avoiding these extreme values, we simply emphasize the role played by the overall recovery dynamics, rather than by the system’s initial response or final convergence.

### C. The temporal dimension of disturbances

We consider a disturbance regime by repeatedly applying disturbances with a given average frequency *f*, over a time period *τ*. For simplicity we assume no correlation in space or in time, so that the time between disturbances is drawn from an exponential distribution with some average frequency (a Poisson process, see Appendix B for details), while the location of the disturbance’s centre is drawn from a uniform distribution.

We use two measures of stability for a system that is disturbed repeatedly, i.e. variability, which measures how far the system ventures from its average value, and persistence, which measures how likely it is to move to the basin of attraction of a different equilibrium. We define variability *V* as the variance in time of the total biomass of the system, given a regime of repeated disturbances. In order to neglect the effect of transients, we calculate *V* over the last 80% of the simulations, which last for 10, 000 time steps. We define the collapse probability *C* as the probability that the system will be in the bare state at the end of a simulation, such that *C* = 0 means no chance of a system collapse, while *C* = 1 means that a collapse is certain. We use a longer simulation time (100, 000 time steps) to calculate *C* since we are interested in predicting a collapse before it occurs. For each of these calculations we run 100 simulations with different randomizations of the location and time of disturbances.

## III. Results

### A. Spanning the spatial dimension of disturbances reveals local ecosystem dynamics

We begin by looking at the response of an ecosystem to a single disturbance with varying spatial extent σ. We focus on disturbances with a fixed overall strength *s* = *s*_0_ for simplicity and clarity, and relax this assumption in the discussion. Thus a global disturbance σ = 1 (Fig. 1, right panels) occurs when *N* is decreased by *s*_0_*K* in the entire system, while a localized disturbance σ = *s*_0_ (Fig. 1, left panels) occurs when *N* is set to zero in a domain of relative size *s*_0_.

**Fig. 1:**
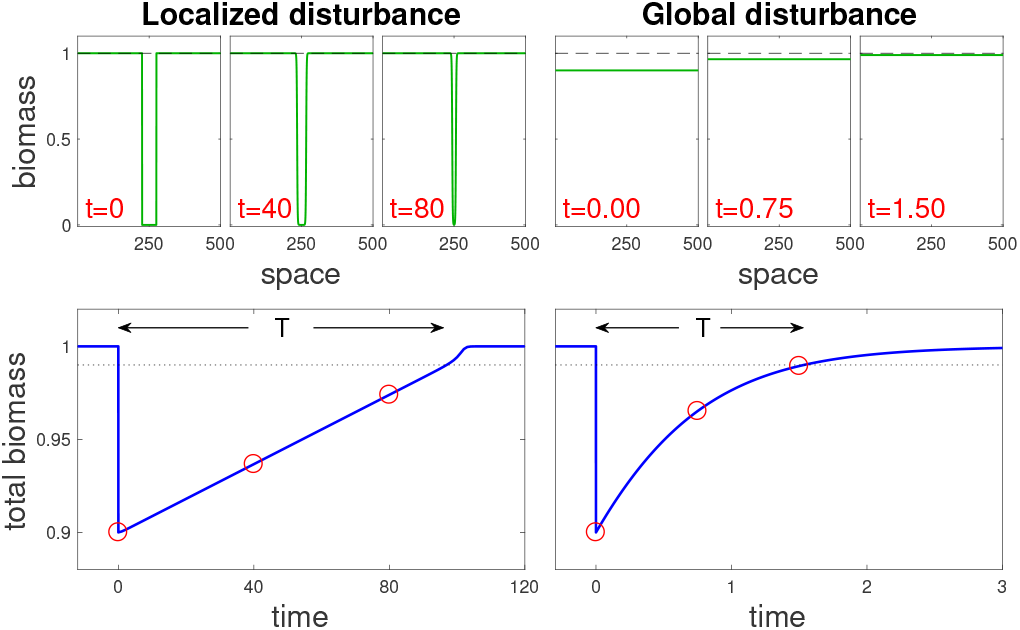
Recovery dynamics following a localized and a global disturbance. (left and right panels, respectively) for the bistable AE model (see Main text). Top panels: snapshots at different times along recovery trajectories, each snapshot showing a biomass spatial profile. Bottom panels show the change in overall biomass over time following the disturbance, where the dotted line denotes the threshold beyond which the system is considered to have recovered, and red circles correspond to the snapshots. Note that the return time *T* from a localized disturbance is much longer than the one from a global disturbance. Disturbance parameters are *s* = 0.1, with σ = 0.1 for the localized disturbance and σ = 1 for the global one.

The response to a disturbance can take two possible forms: isolated recovery due to local processes, and rescue recovery due to incoming biomass from outside the disturbed region. Isolated recovery dominates the system response when each site recovers without the aid of neighboring sites (Fig. 1, right panels). In contrast, rescue recovery occurs when the disturbed region cannot recover without the rest of the system, or when the bulk of the recovery occurs due to such spatial dynamics (Fig. 1, left panels).

To consider the impact of recovery mechanisms in different ecosystems we investigate the behavior of three distinct types of local dynamics, i.e. logistic growth (LG), slow recovery (SR) and Allee effect (AE). For each model we can define a local potential (left panels of Fig. 2), such that its derivative with respect to *N* corresponds to the derivative of *N* with respect to time – i.e. the local dynamics. This means that the local dynamics follow the slope of this potential, so that the biomass density can be thought of as a ball moving from peaks to valleys in the landscape that the potential defines. In both the LG and SR models only one stable equilibrium exists, but the speed of return to the equilibrium may be much slower for low biomass density in the SR model. Two stable states exist in the AE model (the populated state and the bare state).

**Fig. 2:**
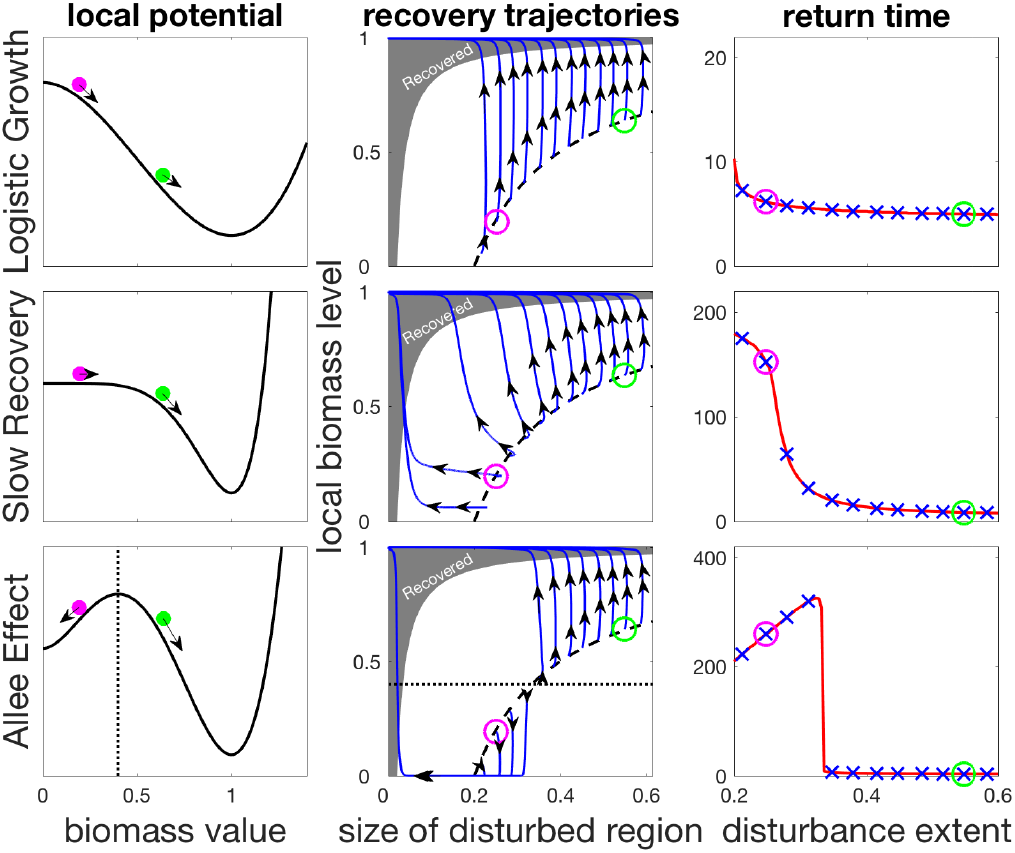
**Contribution of isolated and rescue recovery as a function of disturbance spatial extent** for the three models presented in the main text. The left column shows the local potentials defining local processes. Top row: Logistic (LG) model; Middle row: the highly non-linear SR model; Bottom row: bistable AE model. Middle column: isolated recovery on the y-axis, and rescue recovery on the x-axis. Black dashed line shows the equal disturbance strength used *s* = 0.2 for different disturbance extent σ. Blue lines are recovery trajectories, where recovery is considered complete when trajectories reach the grey shaded region. For the SR and AE model, as disturbances become more localized, a shift is observed from a dominant isolated recovery (upward trajectories) to a dominant rescue recovery (leftward trajectories), impacting return times (right column). The ‘x’ marks in blue correspond to the different trajectories shown in middle columns. The green and magenta circles show initial states following two different disturbances (left and middle columns) and their associated return times (right column). The dotted line (bottom row) shows the local tipping point of the bistable AE model, beyond which local dynamics collapse to the bare state.

The coupling of local dynamics and dispersal results in distinct recovery processes in the three models, as shown by the trajectories in phase-space diagrams in the middle column of Fig. 2. In these panels we unfold the recovery along two dimensions: the horizontal axis denotes the size of the disturbed region at a given time, while the vertical axis shows the biomass density in the disturbed region. Immediately after the disturbance, the system is along the dashed black curve, and it then changes over time until it enters the shaded region where it is considered to have recovered.

If a large part of the trajectory during recovery is horizontal, this means that the disturbed region in shrinking due to rescue recovery, which indicates a lack of local resilience, which would otherwise allow isolated recovery to take place. This behavior reflects the strong nonlinearity of local dynamics, which can be seen in the changing curvature of the local potential (Fig. 2, left column). We can see that for the AE model (Fig. 2, bottom row) the recovery is along a horizontal line for recovery scenarios with a sufficiently small spatial extent, so that regional processes bring about the recovery. In contrast, the recovery is entirely due to local processes in the LG model since the local dynamics are much faster here, while for the intermediate SR model a mixture of the two processes can be seen to take place.

These differences translate into markedly different values of the return time *T* (Fig. 2, right panels). The vertical recovery trajectories that follow all disturbances in the LG model and large-sized disturbances in other models indicate isolated recovery, and hence small values of *T*. For the intermediate SR model localized disturbances lead to a larger contribution of rescue recovery, leading to a sigmoid shape of *T* as a function of disturbance extent σ. The AE model shows a similar behavior of larger *T* following localized disturbances, but the trend here shows a maximum for mid-sized disturbances. This occurs because in bistable systems, the most efficient way to perturb the system is to locally remove biomass just bellow the viability threshold, and then let the system collapse locally. Such a disturbance has an equivalent effect to that of a stronger disturbance that would remove all biomass over a larger region. The spatial recovery process will take longer to recover, thus giving larger return time values (see Appendix A for details). This explains the humped shape of return time as a function of disturbance extent. Since bistability is a sufficient condition for a hump-shaped relationship to occur, the latter could be used as an indicator of bistability. This illustrates the more general idea that considering the spatial dimension of disturbances can allow us to probe the local dynamics of a spatially extended ecosystem.

### B. From variability to collapse under increasing frequency of disturbances

Natural ecosystems are constantly perturbed, leading us to consider a temporal dimension of disturbances, namely their average frequency. We therefore translate the results of the previous section on the response to a single disturbance (Fig. 3, top) into an understanding of temporal variability under repeated disturbances (Fig. 3, bottom). In fact, there is a direct link between the response to a single disturbance and temporal variability in response to repeated disturbances. Indeed, biomass fluctuations are the result of past disturbances, as they integrate short- to long-term responses of the ecosystem to individual disturbances [2]. Variability is a statistic of those fluctuations, and is therefore a function of both the integrated response to a single disturbance and the average frequency *f* of disturbances. More precisely, if *g*(*t*) traces the change in overall biomass through time following a pulse disturbance a time *t* = 0, then variability *V* can be expressed as 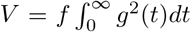 (see eq. refeq:vfg in Appendix B). However, this identity assumes no interaction in space between the different disturbances, and therefore should not hold at high disturbance frequency.

As expected, at low frequency of disturbances the analytical approximation agrees with the numerical simulations quite well for all three models (Fig. 4, second column). For higher frequencies (Fig. 4, third column) where multiple disturbances often take place in the same time frame, we see a slight underestimation of the analytical approximation, although the general trend is well captured. Importantly, variability and return time show the same behavior. We see effects of regional processes on variability for more localized disturbances in both the SR and AE models, where the former shows a sigmoid shape while the latter has a hump shape, which is a consequence of the bistability in the AE model. We note that these trends hold in more general scenarios, such as disturbances with a random extent or following seasonal patterns (Appendix D).

**Fig. 3:**
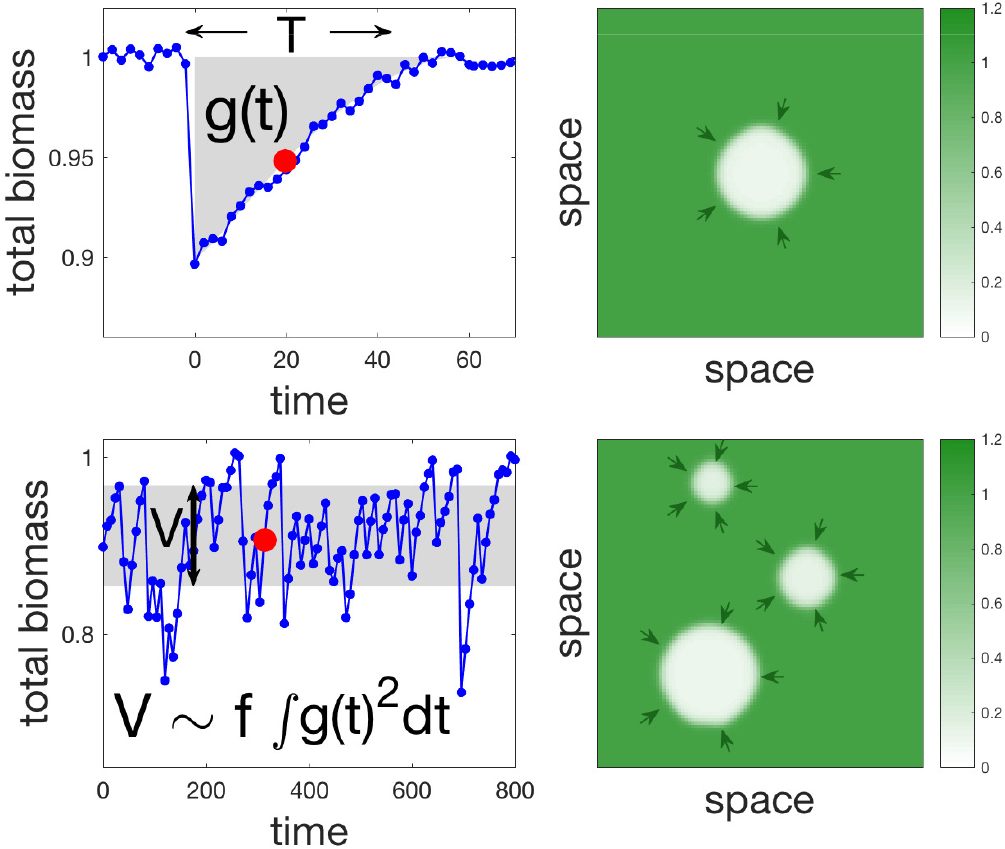
Single and multiple disturbance regimes and the relationship between return time and variability. The left panels show time series of the overall biomass, while right panels are spatial snapshots of the corresponding time-series (red dots in the left panels). The response to a single disturbance is shown in the top left panel. We focus on two of its characteristics: return time *T*, and an integral measure of the transient *g*(*t*) (see main text). The response to multiple disturbances occurring randomly at an average frequency *f* is shown in the bottom left panel. It is summarized by its variability *V* (variance of overall biomass). In the limit of low *f* there is an inherent relationship between return time and variability in the sense that *V* can be approximated by 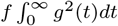. Parameters used *s* = 0.1, σ = 0.11, *f* = 0.025, and *γ* = 2. Random uniform noise was added in left panels to demonstrate how realistic time series might look like.

**Fig. 4:**
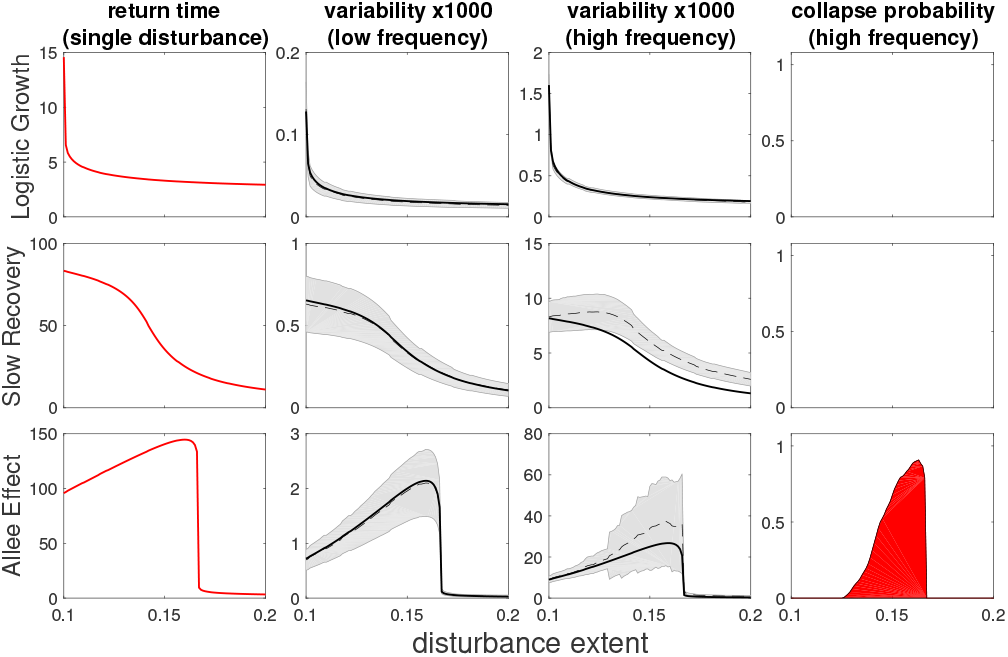
Return time, variability and collapse probability as a function of disturbance spatial extent for three models. (from top to bottom: LG, SR, AE). Left column shows return times (as in Fig. 2) while middle columns show variability under low and high frequency of disturbances, and right column shows collapse probability. The black dashed (solid) line is a numerical (analytical prediction) value of variability, with grey shading noting error estimation. Deviation from this prediction implies some degree of interaction between disturbances. Return time and variability are qualitatively similar with low dependency of disturbance spatial extent for the LG model but a much stronger dependency when local dynamics are highly non-linear (SR and AE models). In the case of the bistable AE model we recognize a non-monotonous “hump-shaped” dependency with disturbance extent, with mid-sized disturbances causing the most severe response. Disturbance parameters were *s* = 0.1, σ = 0.1, and for low frequency: *f* = 0.002, while for high frequency: *f* = 0.02.

At this point it would appear that the temporal dimension of disturbances *f* is not as informative on ecosystem behavior as the spatial dimension of disturbances σ. However, as *f* is increased further, a discrepancy between variability and its prediction based on recovery from a single disturbance starts to grow. This signals that the disturbances start to interact with each other, a phenomenon that is not captured by our approximation. Disturbances start to aggregate in space, which can substantially increase variability (Appendix C) due to large excursions towards low total biomass levels. For bistable systems such as the AE model, such excursions can lead to a collapse of the whole system. This is evident in the two last columns of Fig. 4, in which we see, for the AE model, that the values of σ for which the discrepancy of variability is highest precisely corresponds to the values of σ for which the collapse probability is most signifi-cant. Thus, at high frequencies, disturbances of similar strength but different spatial extent lead to dramatically different responses. This example highlights the fact that the combination of spatial and temporal dimensions of disturbances can have a drastic effect on ecosystem stability.

## IV. Discussion

Investigating the role of the spatial and temporal dimensions of disturbances in ecosystem stability, we obtained four main results: (1) In comparison with a global disturbance, a localized one of the same strength can initiate a fundamentally different, and much slower, ecosystem response, especially when local dynamics are nonlinear. (2) The return time from a single disturbance and the temporal variability caused by repeated disturbances show the same trends, even for locally intense (and therefore nonlinear) disturbances. (3) The relationship between a system’s response and the spatial extent of the disturbances it experiences reveals its underlying dynamics. For instance, a hump-shaped relationship between return time and the spatial extent of the disturbances may indicate bistability. (4) The correspondence between return time and variability breaks down for high disturbance frequencies. This discrepancy signals the occurrence of spatial interactions between disturbed regions, which, in turn, may lead to a regime shift.

Uniquely to our work, we considered systems locally pushed far from their equilibrium, and even to a different basin of attraction. These locally intense disturbances allow rescue recovery, mediated by dispersal, to dominate the ecosystem response. In the case of the bistable (AE) model this glimpse outside the basin of attraction of the populated state is the direct cause of the hump-shaped trends of variability and return time as a function of disturbance extent. In fact, the front propagation that drives rescue recovery contains information about the ecosystem’s basins of attractions, reflecting the existence of alternative stable states and the transient dynamics between them. Thus, by observing the ecosystem’s response to localized disturbances, rescue recovery allows us to probe ecosystem dynamics far from equilibrium. This reasoning could be taken further by focusing on regions where rescue recovery takes place, e.g. analyzing the plant community structure at transition zones between grassland and forest in a savanna ecosystem. This could allow the development of novel empirical methods to better assess and understand the dynamical properties of spatially extended ecosystems.

Spanning the spatial dimension of disturbances could thus allow us to detect nonlinearities in ecosystem behavior, revealed by the increasing local intensity of disturbances (see Fig. 2). One might expect that along the temporal dimension of disturbances, increasing their average frequency could also reveal nonlinear effects, since the ecosystem becomes more strongly disturbed. In fact, increasing frequency has only a trivial linear effect, as reflected by the relation we found between return time and variability (see Fig. 3). Beyond some threshold, however, a response of a different kind emerges, due to spatial interactions between disturbed regions which aggregate in potentially large-scale clusters. This causes a higher variability than expected and can, consequently, cause a global loss of persistence or a regime shift. Importantly, the two dimensions, spatial and temporal, must now be considered in conjunction. The threshold beyond which aggregation occurs depends strongly on the spatial extent of disturbances and hence the associated response is not a mere superposition of responses to single disturbances. In other words, this finding highlights and explains how the interplay between the spatial and temporal dimensions of disturbances can lead to drastic consequences, such the loss of persistence. Since our findings are purely theoretical, it would be enlightening to elucidate the prevalence of this interplay in empirical systems that have undergone regime shifts (e.g. green Sahara [23]).

As previously mentioned, in bistable systems the relationship between return time (as well as variability) and the spatial extent of disturbances is hump-shaped. This relation could be used as an indicator of bistability, assessed empirically by comparing time series of the same ecosystem in different regions with estimates of the intensity of single disturbances. Its implications for ecosystem management depend on the type of disturbances considered. Anthropogenic disturbances that are largely controlled, such as logging in forests, can be better planned to avoid both an unpredictable yield due to high variability and an overall collapse due to loss of persistence. For many natural disturbances control is neither possible nor desired, but predicting their effects and the possibility of regime shifts is paramount.

In order to focus on the role of the spatial properties of disturbances and allow a clearer presentation, we conducted our analysis assuming disturbances of constant overall strength. It is straightforward to extend the analysis to more general settings, such as a random extent of disturbances and seasonal patterns (see Appendix D for details). It is particularly interesting to consider the case of different values of disturbance strength *s*. As shown in Fig. 5, if we randomly choose a set of points with different values of strength *s* and extent σ, we can use these to reconstruct a normalized version of the dependency of the different stability measures on disturbance extent. Thus we can use the different phenomena described previously, such as a hump-shape relationship as an indicator of bistability, under more general conditions. Although we considered simple spatially homogenous models, our results should apply to a wide range of ecosystems. Forests, savannah and shrublands might be good examples of ecosystems to which our models apply since disturbances such as fires and grazing occur frequently and are often localized, and the recovery of plant communities often follows complex succession dynamics driven by spatial processes [1, 27, 31]. Our results, however, need not be restricted to such spatially homogeneous systems. Although we built our theory using spatially uniform models, this simplifying feature is not essential to our arguments, which only require a notion of locality. Therefore, our theory may also be relevant to less homogeneous ecosystems, such as mountain lake networks, coral reefs and riverine systems. Indeed, such ecosystems undergo different disturbances that are often strongly localized, and their dynamics may be sufficiently nonlinear [4, 8, 12].

**Fig. 5:**
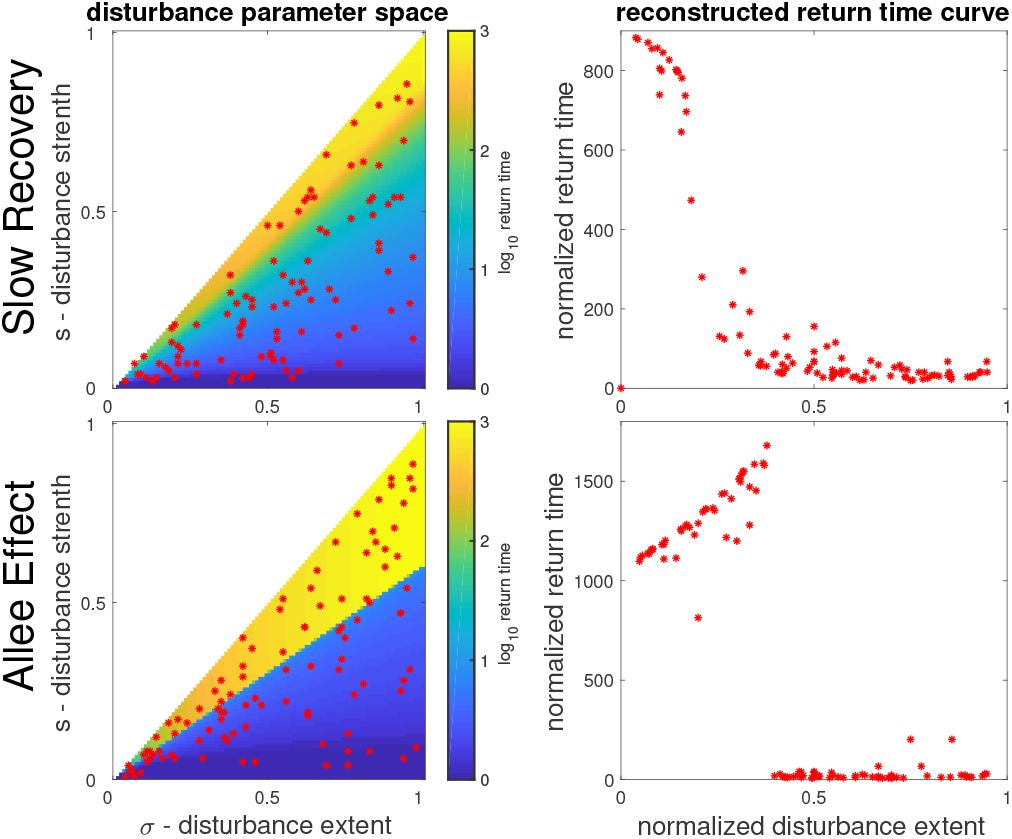
Reconstruction of return time vs. disturbance extent curve from the more general parameter space of disturbance properties. Top (bottom) panels correspond to the SR (AE) model. Left column shows the return time over the parameter space of disturbance extent σ (x-axis) and of disturbance strength *s* (y-axis). Right column shows the corresponding reconstruction of the return time curve, using 100 randomly chosen points (red asterisks) in the parameter space. The return time values are normalized by the disturbance strength *s*, while the normalized disturbance extent is defined as 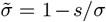. Note that the hump (sigmoid) shape of the curve for the AE (SR) model are easily recognizable from these reconstructions.

Our work is a step towards a quantitative account of spatial and temporal dimensions of disturbances, and their interplay with local and regional ecosystem dynamics. This is an important goal in the context of global change. Disturbances are of increasing frequencies and occur at different scales, while the spatial structure of ecosystems themselves is altered by land use change. It is thus important to build a framework in which we can understand and predict the ecological impacts of this complex interplay.

## Authorship

YRZ, JFA and ML designed the study, YRZ and JFA performed the research; YRZ, JFA and ML wrote the manuscript.

## Acknowledgements

We wish to thank Bart Haegeman and Matthieu Bar-bier for helpful discussions and comments on a previous version of the manuscript. This work was supported by the TULIP Laboratory of Excellence (ANR-10-LABX-41) and by the BIOSTASES Advanced Grant, funded by the European Research Council under the European Union’s Horizon 2020 research and innovation programme (grant agreement No 666971).

## Methods

All simulations used to create the figures were made using Matlab with the library RDM: https://github.com/yzelnik/RDM

Code that shows how simulations and calculations for the main figures were made can be found at: https://github.com/yzelnik/stddes-scripts

## Appendix A: Approximation of return time for the AE model

To understand the shape of the return time curve *T* (σ) for the AE model, as seen in bottom right panel of Fig. 2, we consider the initial response of the system just after the disturbance has occurred. If we ignore the effect of diffusion, then the domain that was not disturbed does not react at all, while the disturbed domain either rebounds back to the vegetated state *N* = *K*, or falls further to the bare state *N* = 0. This depends simply on whether the current level of biomass *N*_0_ (immediately after the disturbance) is higher or lower than α. If we note the system size as *L*, the spatial extent of the disturbance as σ, and the disturbance strength (percentage of biomass cut) as *s*, then the biomass value at the disturbed region just after the disturbance occurred is: 
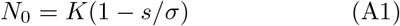

We therefore have a critical value of σ when *N*_0_ = *α*, which will decide whether the disturbed region would rebound without the effect of diffusion, as: 
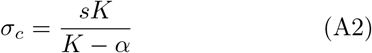

As shown in Fig. A1 with a vertical red line, this approximation gives a good indication to where the return time changes behaviour rapidly. For larger disturbances σ > σ*_c_*, each single point would recover from the disturbance on its own accord. Moreover, the contribution of spatial processes to this recovery would be negligible if the rate of isolated recovery *r_iso_* is much faster than the rate of rescue recovery *r_res_*, namely *r_iso_ ≫ r_res_*. We can use simple dimensional considerations [19] to estimate what are the conditions for this situation.

**Fig. A1:**
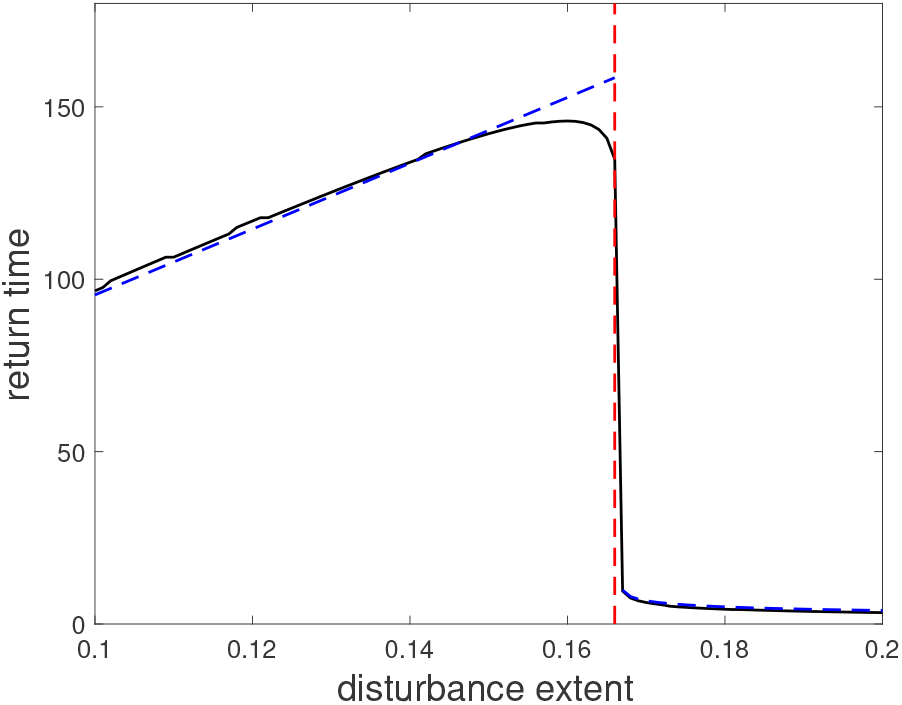
Approximation of the return time for the AE model. The black curve denotes the return time for the AE model (see Fig. 2), where we set *s* = 0.1. The red vertical line shows the approximate separation point between the different regimes of recovery, 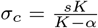. The two blue curves show the approximations of the return time in the two different regimes. These approximations are calculated as described in the text and multiplied by a factor of 0.9 since the return time (as shown in black) is defined as recovery of 90% of the biomass (see Methods).

First, we can estimate *r_iso_* by *r*, since it is the only parameter with units of time and no relation to space. This estimation will break down close to the unstable equilibrium (since the recovery rate will go to zero), i.e. when σ is close to σ*_c_*. To estimate *r_res_* we need to consider how fast a front between the disturbed and undisturbed domains moves. In general, the front’s speed *u* between two domains in a model that can be written as *N_t_* = *rf*(*N*) + *d*∇^2^*N* would be proportional to both the rate of local dynamics *r* and the rate of diffusion *d* as 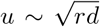, due to dimensional considerations. The area of the disturbed region is σ*L*, so that the overall time to recovery is the ratio between area and speed, 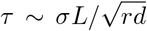. We can thus estimate the rescue recovery rate as 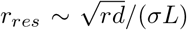, and compare this to *r_iso_*. This gives us the condition 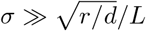, which is easily met in the parameters we choose since *L* is large enough. Therefore to approximate the rate of recovery for σ > *_c_* we can just use the return time when we ignore spatial effects (*d* = 0). Indeed, this estimation, shown by the right-more blue curve in Fig. A1, works well, even for lower values of σ close to the critical value σ*_c_*.

For the opposite regime, σ < σ*_c_* we can easily assume that isolated recovery is negligible, since in the bistable AE model isolated recovery is not possible when locally *N* < *α*. To estimate the rescue recovery we again consider the speed of a front between the disturbed and undisturbed domains. Here, both of these can be assumed to be the two stable states, and under these conditions the front speed is a constant that depends only on the model parameters [19]. In particular, the front speed *u* grows with the rate of local dynamics *r* and diffusion coefficient *d* as 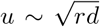, and for the AE model considered here, *u* also grows with *α* once *α* > *K*/2 (the front is stationary at *α* = *K*/2 due to the symmetry between the two stable states for these parameters). If we assume that once a disturbance occurs, the time until the disturbed region collapses to the alternative state is negligible, then the recovery time *T* in simply the time it takes the fronts to take over the disturbed region: 
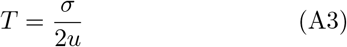

This approximation is shown in the left-more blue curve in Fig. A1. While there is a clear under-estimation of the return time in this case due to complex front dynamics, the overall trend and the value for very localized disturbances is well approximated.

## Appendix B: Relation between return time and variability

In this appendix we derive a general relation between recovery dynamics following a single disturbance, and variability under repeated disturbances. In this manuscript we formalize disturbance events as a realization of a point process. We therefore begin by explicitly defining a point process that we use for our derivation as well as the assumptions we make for this derivation. We then detail the derivation itself, and finally we discuss the main results of this derivation.

### Definitions

A point process (p.p.) Φ is a random sampling of points in 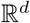 such that the number of points sampled in a bounded set is always finite [3]. We note Φ (*A*) as the restriction of a p.p. to a given set *A* and we note *φ*(*A*) = *{x*_1_*, x*_2_*,…} ⊂ A* as any of its realizations. Let *N*(*A*) be the random variable representing the number of points in *A* sampled by Φ. The intensity measure Λ of counts the expected number of Φ point sampled from a given set, i.e. 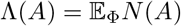. Note that, if *A* and *B* are disjoint sets, *N*(*A*) and *N*(*B*) are independent random variables so that Cov_Φ_(*N*(*A*)*, N*(*B*)) = 0. We call a Poisson process a p.p. such that, for any *A*, the mean number of points sampled is also equal to the variance so that 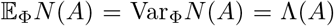. A homogeneous Poisson process satisfies Λ (*A*) = *f*|*A*| where |*A*| is the usual volume of *A*. In this case, this simply means that Φ (*A*) is a uniform sampling of points of *A*, with an expected number of points proportional to the volume of that set.

Our derivation, as given below, follows from the intuition that if the disturbances are not too frequent, so that they do not often interact in space, then we can use the superposition principle to find a relation between the response to one disturbance and the response to multiple disturbances. More concretely, our reasoning, as illustrated in Fig. A2, requires the following assumptions:

**Fig. A2:**
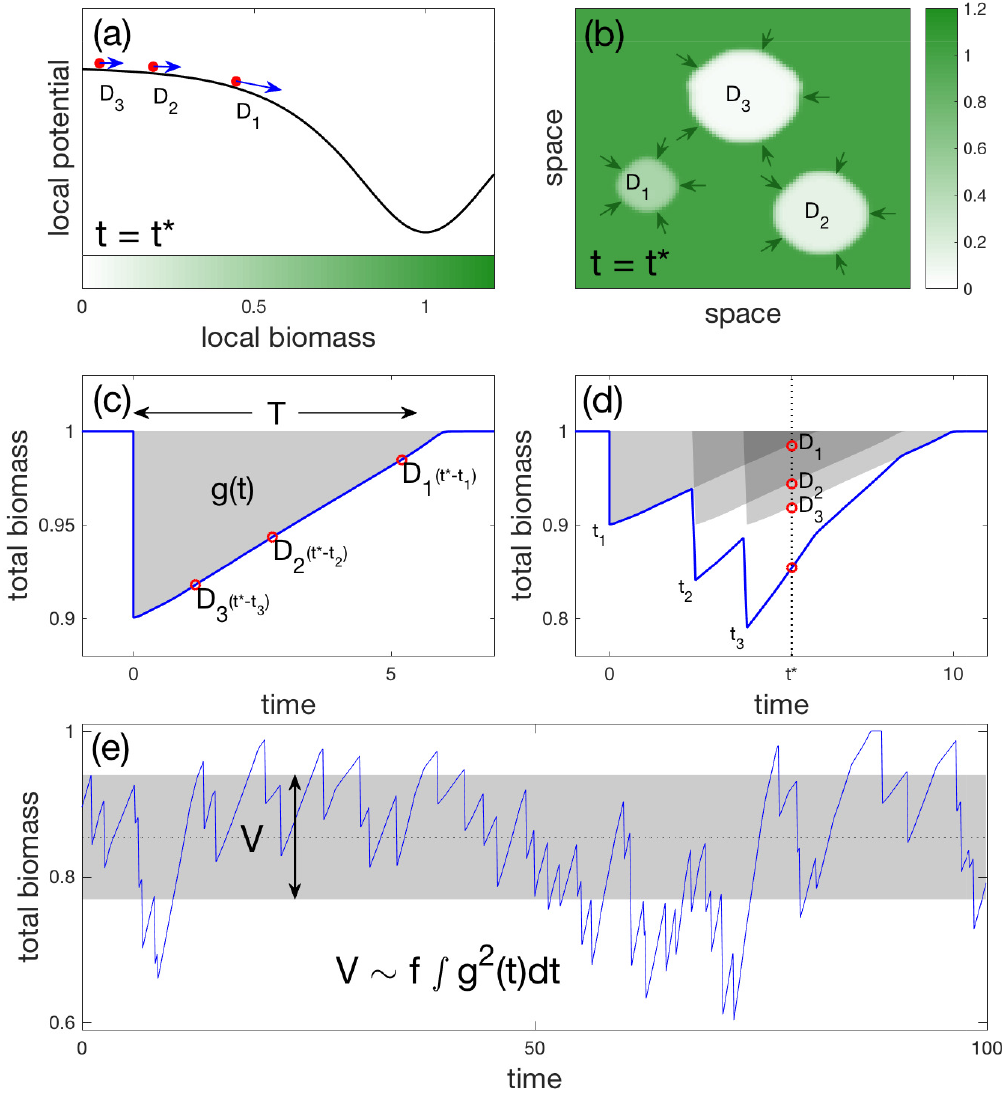
Illustration of how return time following a single disturbance translates into the variability under a regime of repeated disturbances. (**a,b**) Snapshot at time *t*\* of the local and regional dynamics, following three different disturbances (*D*1, *D*2, *D*3) at times (*t*_1_, *t*_2_, *t*_3_). The local dynamics (a) are shown using a potential (similar to Fig. 2), with three balls and arrows corresponding to the local state at each disturbed region. A snapshot of the two-dimensional system shows the regional dynamics**(b)**, as each disturbance occurred in a different location in space in a different time, and therefore has a different size and different biomass value in the disturbed region. **(c)** The trajectory of a system following a single disturbance is shown, where the current state at time *t*\* of each disturbance (if it occurred alone) is noted by a red circle. **(d)** The overall biomass over time due to the three disturbances can be approximated well by adding the three trajectories following a single disturbance in appropriate times, as shown by the grey shading. The effect at time *t* of each disturbance and all together is shown by the red circles. **(e)** A regime of repeated disturbances at different times results in a noisy time series (blue line) that can be measured by the variance (grey shade). This measure of variability *V* can be well approximated as the product of the average frequency of disturbance *f* and the second moment of the trajectory following a single disturbance *A*.

1. We assume disturbance events to be realizations of a Poisson p.p. Φ in 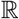 of intensity measure Λ.
2. We assume no spatial interactions between disturbances, thus neglecting co-occurrence events, where multiple disturbances co-occur in a small region.

### Derivation

Let us denote *g*(*t*) as the recovery function from a single disturbance, so that *g*(0) = *sK*, where *s* is the relative overall strength of the disturbance, and by our definition of return time *T_R_* we have: *g*(*T_R_*)*/g*(0) = 10%. Let *φ*([*−∞, t*]) be a random realization of all past disturbance events. The displacement time series of overall biomass under a repeated disturbance regime, *h_φ_*(*t*) = *K −N_tot_*(*t*) will be 
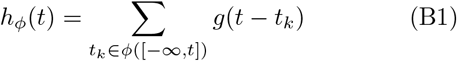

Note that for Eq.(B1) to hold, assumption (2) is necessary only if the dynamical response to an individual disturbance is non-linear. If the response is linear, a superposition principle allows Eq.(B1) to be true [2], even without assumption (2). In fact, for global disturbances (σ = 1), assumption (2) cannot hold, yet Eq.(B1) still holds because a global disturbance typically induces only small displacements from *K*, so that the recovery is essentially linear.

By ergodicity of the point process, taking an average over long times for one realized time series is equivalent to taking a point average (say at time 0) over realizations, so that 
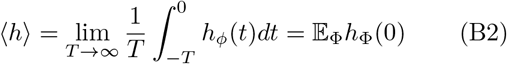

We are thus left to analyse random variables of the form 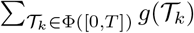, a realization of which is ∑*_tk_*_∊φ_([0,*T*])*g* ^(*t*_k_^ (note the change of variables −*t_k_* → *t_k_*, so as to restrict to positive times). We may approximate *g*(*t*) with arbitrary precision by a piece-wise constant function 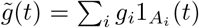where the _*Ai*_’s are mutually disjoint intervals such that ∪ *A_i_* = [0, *T*] and the functions 1_*Ai*_(*t*) are defined by 1_*Ai*_(*t*) = 1 when *t* ∊ _*Ai*_ and 0 otherwise. The numbers *g_i_* approximate the function *g* in *A_i_*. Importantly 
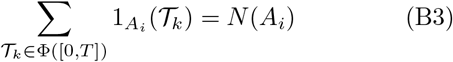

From assumption (1)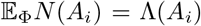 where Λ is the intensity measure of Φ. We thus see that 
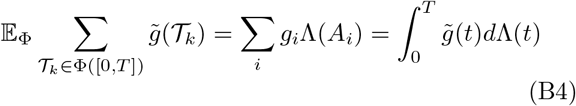

Because the approximation of *g(t)* by 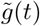 is arbitrarily good this finally yields: 
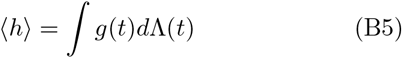

In the main text we defined variability as the variance of *h_φ_ (t)*. If we repeat the above reasoning on 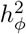 instead of *h_φ_* we get that, with arbitrary accuracy, 
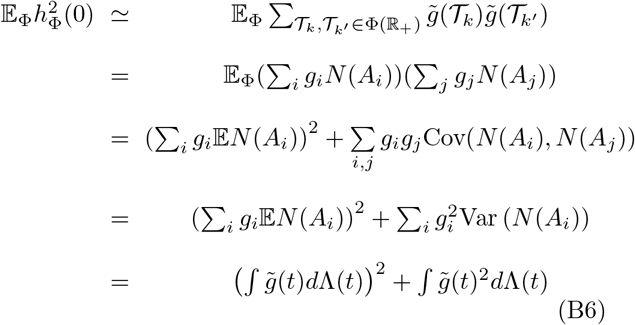
where the last two lines follow, respectively, from the fact that *N*(*A_i_*) and *N*(*A_j_*) are independent r.v. if *i* ≠ *j* (no temporal correlations) and the fact that, for a Poisson process,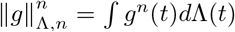. In terms of the integral norms 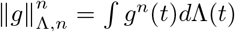, what we have proved is that, under assumption (1) and (2) 
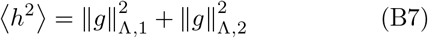
so that variability becomes 
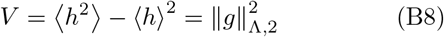

### Discussion

In the particular case of a homogeneous Poisson process of intensity *f*, so that Λ(*A*) = *f*|*A*| then we have that 
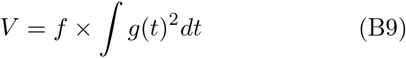
as illustrated in Fig. A2. Note that homogeneity is not essential to our arguments. For instance, if *t* is expressed >in years, then seasonality effects could be modelled by taking, 
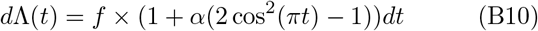
with 0 ≤ *α* ≤ 1. What is essential is the absence of temporal correlations between disturbance events, which assumes that a disturbance does not inuence the occurrence of future disturbances. This is not always true in natural systems, (e.g. a drought can make a fire more likely), but is a reasonable assumption for most climatic events.

Note that we assumed so far that all disturbances had the same spatial extent σ. This assumption can easily be relaxed by assuming σ to be a random variable, with p.d.f. *p*σ. The contributions to the overall variability of disturbances of size σ^′^ ∊ [σ,σ + *d*σ] is 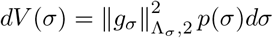; so that 
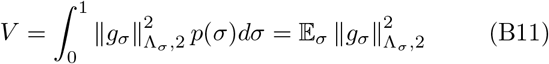

In this sense, the relationship Eq.(B11) is quite general. However, it requires assumption (2) to hold, especially for localized regimes of disturbances, for which the response will typically be highly non-linear. As we explain in the next section, departures from this prediction is the sign that disturbances are aggregating into long-lived disturbed regions, a process that increases variability and, in bistable systems, can lead to a regime shift.

Finally, it is interesting to note that had we assumed no temporal overlap between responses to disturbances (assuming a very low frequency of events, for instance) then we would have had 
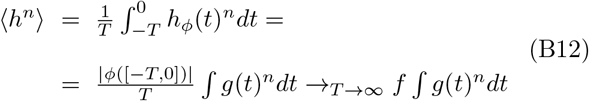
where |*φ*([−*T*, 0])| is the number of disturbance events that have occurred in [−T, 0]. For a homogeneous p.p. and for *n* = 1 we recover Eq.(B5). However, for *n* = 2 we do not find Eq.(B7). This shows that assuming no temporal overlap between responses to compute V would induce an error equal to the squared mean displacement 〈*h*〉^2^. An error that can easily be significant (but is of second order in *f* as the frequency of events goes to zero).

## Appendix C: Aggregation of disturbances

As discussed in the Results section, the aggregation of disturbances is the main reason for the under-estimation of variability and for high collapse probability(Fig. 4). We describe here how this phenomenon affects variability and collapse probability.

We define a disturbance aggregation as a situation when multiple disturbances occur within the same time frame in different locations so that effectively a large contiguous region in the system is disturbed at a given time. This aggregation typically reduces the total biomass, and more significantly, leads to an overall slow recovery when compared to a simple addition of multiple non-interacting disturbances. As shown in Fig. A3, occurrence of such aggregation changes the recovery trajectory considerably (top panel), which leads to higher variability (bottom panel).

**Fig. A3:**
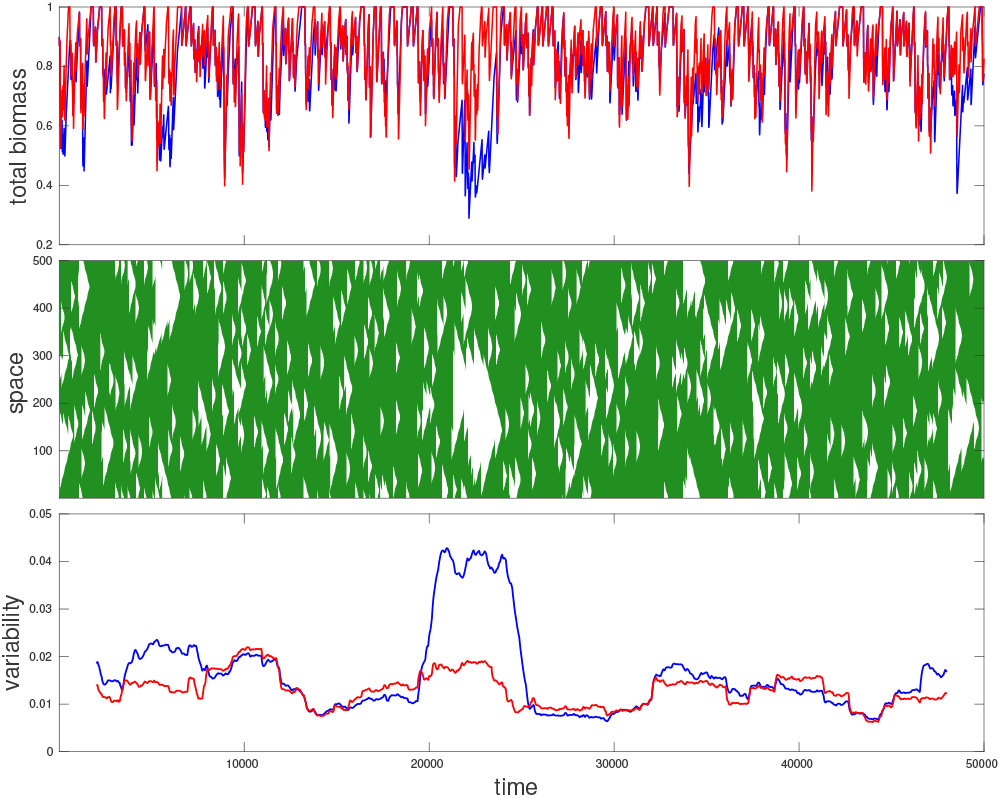
Dynamics of single simulation with multiple disturbances, demonstrating the effect of disturbance aggregation. The dynamics of a single simulation of the AE model with disturbance frequency of *f* = 0.01 and mid-sized disturbances σ = 0.15 are shown in the three panels. This is compared with a “constructed time-series” where the effects of a single disturbance are added multiple times to form a time-series that behaves as a simulation where disturbances do not interact in space (so that their effect is always linearly additive). The top panel shows the overall biomass over time, for both the actual simulation (blue) and the constructed time-series (red). The middle panel shows a space-time simulation where the vertical axis shows the space of a one-dimensional simulation, and colour depicts the amount of biomass (darker green denotes more biomass). The bottom panel shows variability calculated over a running time-window of *τ_v_* = 4000, where for each point shown, the variability was calculated for part of the time-series with length *τ_v_* that is centred on that time point. The variability of the actual simulation is shown in blue, whereas the variability for the constructed time-series is shown in red. A large disturbance aggregation is seen starting *t* = 20000, which lowers the biomass and increases the variability considerably (blue), with no counterpart for the constructed time-series (red). Parameters used: *r* = 0.5, *s* = 0.1.

These events are more common for mid-sized disturbances, which is the reason that the under-estimation is most prominent for these values of σ. This is because an aggregation is more likely to occur when disturbances have both a longer time-span and a larger spatial extent. Larger values of σ mean that the size of each disturbed region is larger, and therefore it takes less disturbances to cover a certain region. On the other hand, more local disturbances (smaller σ) tend to lead to slower recovery (this can be quite different between models, depending on local dynamics), and hence the effect of each disturbance is retained for a longer time. Overall mid-sized disturbances are the middle-ground between these two conditions, and it is therefore there that disturbance aggregation occurs most frequently. For the specific case of a bistable system such as the AE model, mid-sized disturbances also have the slowest recovery, so the effect is even stronger.

We note that for bistable systems, a collapse to the alternative state is possible if enough disturbances push the system to the other state. This is essentially an extreme case of disturbance aggregation, where the disturbances cover the entire system. Therefore, when increasing the frequency of disturbances we first see a collapse occurring for mid-sized disturbances, and only for higher frequencies do we see it for smaller and larger extent of disturbances (see Fig. A4).

**Fig. A4:**
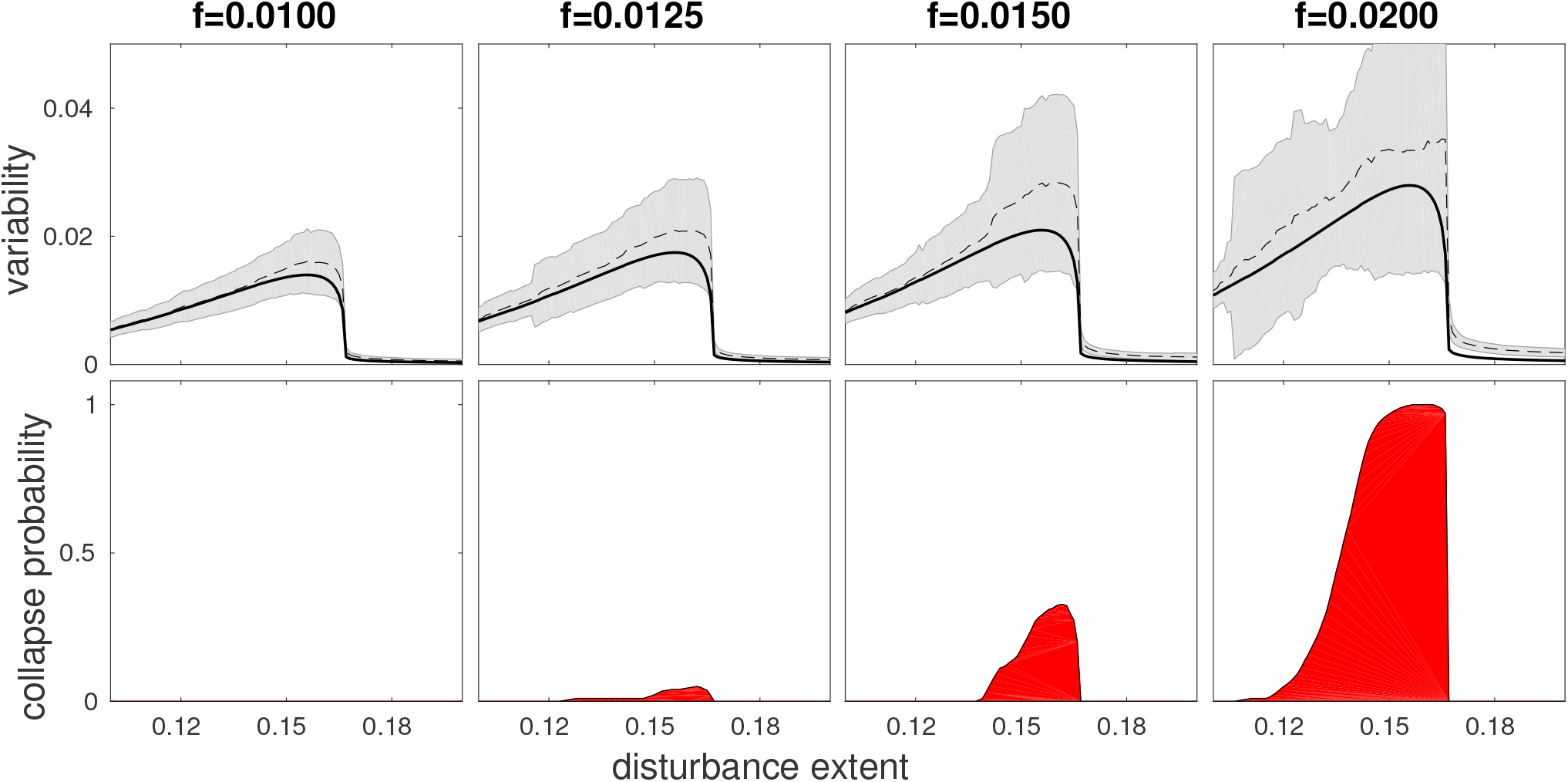
Variability *V* (top) and collapse probability *C* (bottom) in the bistable AE model, for increasing frequencies of disturbances *f* = {0.01, 0.0125, 0.015, 0.02}. The variability calculated from simulations is noted with a dashed black line, with grey shading noting the error estimation of the standard deviation, while the analytical approximation (Appendix B) for it is shown with a solid black line. Parameters used: *r* = 0.5, *s* = 0.1.

## Appendix D: Generalization of results

The results presented in this paper have focused on a simple model setup for the sake of clarity, but they can be easily generalized in various settings. For example, in eq. B10 in Appendix C we show how temporal seasonality can be taken into account and give the same overall results. We further demonstrate here that our results are not specific to a particular spatial structure, with two specific examples. We compare between one-dimensional (1D) and two-dimensional (2D) systems, looking at the results for return time, variability and collapse probability. We also consider a regime of repeated disturbances where the size of disturbances is not the same every time, but rather taken from a probability distribution.

To demonstrate that the specific spatial structure of the ecosystem is not a fundamental part of our theory, we show here results for a two dimensional (2D) ecosystem, and compare them to results for a one dimensional ecosystem (1D) that we show in the main text. For the 2D case we consider an ecosystem with the size of 200×200 with periodic boundary conditions, and use disturbances with a circular shape. As can be seen in Fig. A5, the qualitative properties for return time, variability and collapse probability, as described in the main text, are the same for the 1D and 2D ecosystems. Note that the specific frequencies that are compared are not the same, but rather chosen to show the similarity in trends between the 1D and 2D ecosystems. The trends of variability and collapse probability as described in the main text hold under more general conditions. We consider here a more realistic scenario by allowing disturbance’s size to randomly vary around a fixed average. As an example we set σ to be randomly chosen from a Gaussian distribution with a standard deviation of 0.05 around a given average (Fig. A6). Both the variability and collapse probability show a hump-shaped relationship with the spatial extent of the disturbance in the bistable AE model (top panels), while for models with a single equilibrium such as the SR model, no such behaviour is seen (bottom panels).

**Fig. A5:**
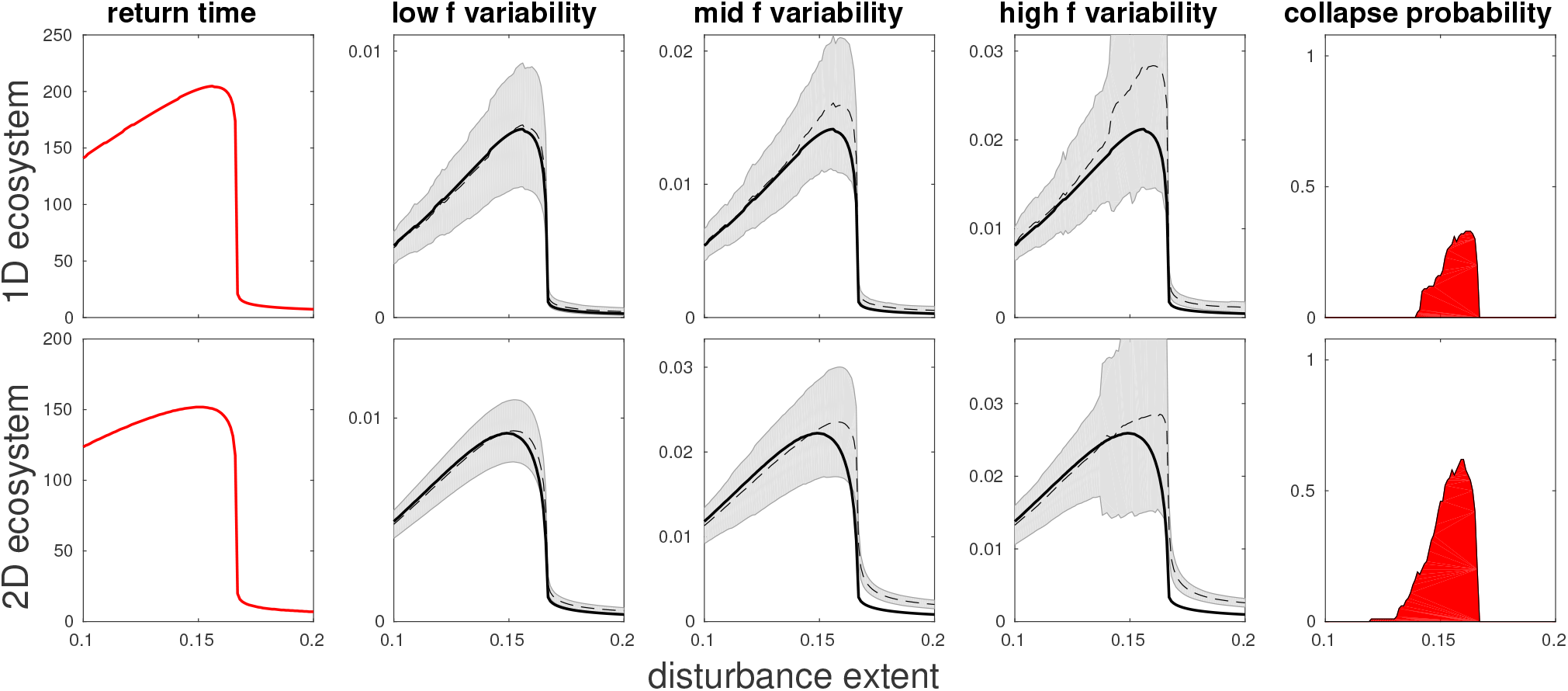
Comparison of return time, variability and collapse probability between 1D (top) and 2D (bottom) ecosystems of the AE model. The left-most column show the return time, the three middle columns show variability for different frequencies, and the right most column shows collapse probability for the highest frequency. The values of *f* are chosen to show similar behavior between the 1D and 2D case (*f* = 0.005, 0.01, 0.02 for 1D ecosystems, *f* = 0.01, 0.02, 0.05 for 2D ecosystems). In the middle columns the dashed line shows numerical values of variability with grey shading noting error estimation. The analytical prediction (based on the response to a single disturbance) is shown in a solid black line, where deviation from this prediction implies some degree of interaction between disturbances. Parameters used: *r* = 0.5, *s* = 0.1. 2D system is of size 200×200.

**Fig. A6:**
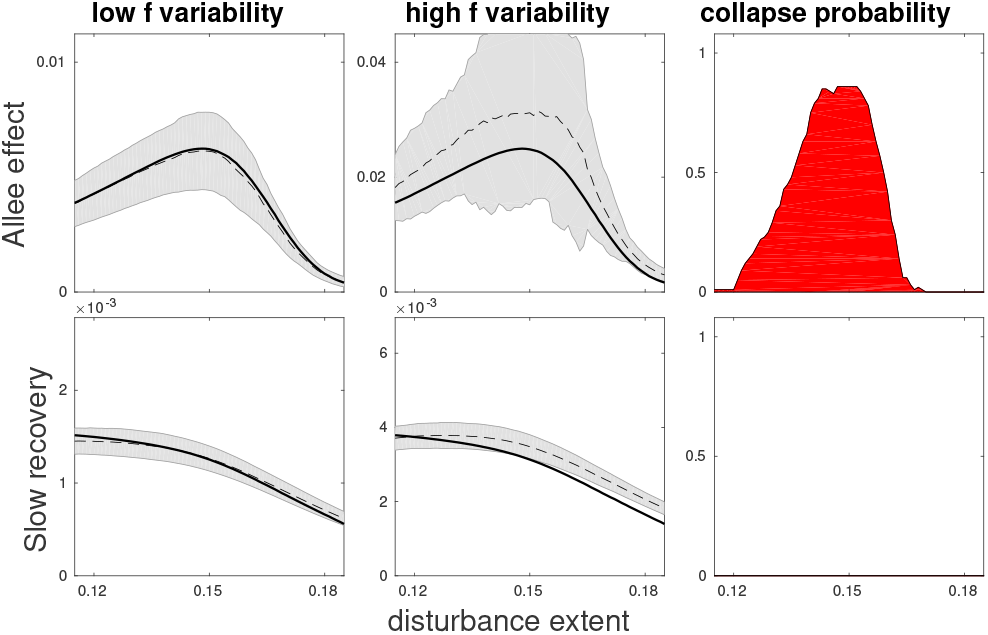
Variability and the probability of collapse under a repeated disturbances with randomly distributed spatial extent. Results are for the locally bistable AE model (top panel) and the SR model (bottom panel) that has a single equilibrium (compare to Fig. 4). Left and middle columns show variability (for low and high frequency respectively), where the dashed line shows numerical estimations with grey shading noting error estimation. The analytical prediction (based on the response to a single disturbance) is shown in a solid black line, where deviation from this prediction implies some degree of interaction between disturbances. Right column shows collapse probability for the high frequency of disturbances. Disturbance extent is taken from a Gaussian distribution with a standard deviation of 0.01, where the mean is changed along the x-axis between 0.1 and 0.2. Comparing with Fig. 4 we see that allowing random variations in disturbance extent leads to variability and collapse probability that are more of a smooth function of average disturbance extent while preserving the hump shape for the bistable AE model. Low and high frequencies used are *f* = 0.005, 0.02 for the AE model and *f* = 0.02, 0.05 for the SR model. Parameters used: *r* = 0.5, *s* = 0.1.

